# The role of contact inhibition in intratumoral heterogeneity: An off-lattice individual based model

**DOI:** 10.1101/036467

**Authors:** Jill Gallaher, Alexander R.A. Anderson

**Affiliations:** Integrated Mathematical Oncology (IMO) Department, Moffitt Cancer Center, Tampa, FL 33605 USA.; IMO Department, Moffitt Cancer Center, Tampa, FL 33605 USA.

## Abstract

We present a model that shows how intratumoral heterogeneity, in terms of tumor cell phenotypic traits, can evolve in a tumor mass as a result of selection when space is a limited resource. This model specifically looks at the traits of proliferation rate and migration speed. The competition for space amongst individuals in the tumor mass creates a selection pressure for the cells with the fittest traits. To allow for organic movement and capture the invasive behavior, we use an off-lattice individual-based model.

## I. Trait Selection in a Heterogeneous Tumor

Individual-based models are often used in cancer research to study how cells interact with each other and their environment. For a simple model of a tissue or mass of cells, one might create a square lattice with each lattice point representing a cell. This is a straightforward and computationally efficient way to represent many cells within spatially regular neighborhoods and works well for a solid growing tumor. However, some tumors may be rather diffuse and invasive, and confining cells to a fixed lattice can hinder a more accurate treatment of cell migration. To illustrate this, we consider an important problem in cancer: intratumoral heterogeneity, which has been shown to complicate cancer diagnosis [1, 2, 3, 4] and interfere with treatment strategies because of drug resistance and recurrence [5, 6, 7, 8]. Here we investigate intratumoral heterogeneity using two fundamental tumor cell traits: proliferation rate and migration speed, to understand how they might evolve when space is a limited resource.

We grow a tumor using an off-lattice individual-based model starting with a small clump of cells with a heterogeneous mix of phenotypes (defined by differences in proliferation and migration traits). When the neighborhood around a cell is filled, it will enter a quiescent sate, in which it will stop both proliferating and moving, so it essentially cannot create any more progeny. We want to investigate which combinations of traits come to dominate given the competition for space amongst neighbors. Each cell moves through the cell cycle as time progresses, and when it completes its cycle (defined by the proliferation rate), it searches the immediate spatial neighborhood for space to divide (see Fig. 1A). If there is an angle in which there will be no overlap with another cell, the cell will divide into two and each daughter cell will keep the phenotype of the mother cell, otherwise, it will go quiescent. We assume that individual cells move with a persistent random walk [9, 10], such that they follow the same direction until reaching the persistence limit, then turn and start again. If they collide with another cell before reaching the persistence limit, then they follow a new trajectory at an angle reflected along the normal to the plane of collision.

**Figure 1.**
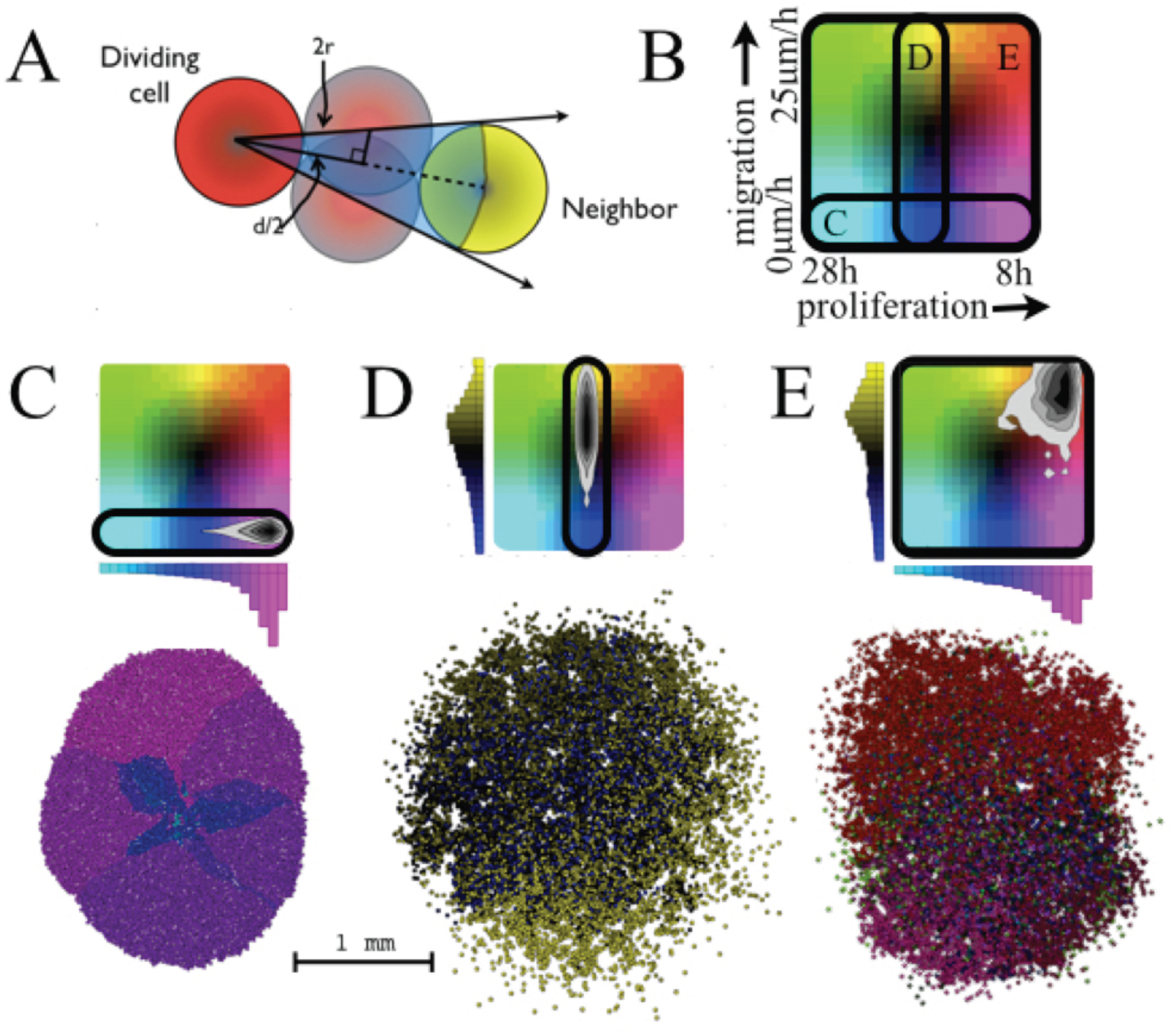
A) Angle exclusion due to occupied space by cells. B) Allowed trait space for (C-E) showing colors of cells with combinations of proliferation and migration traits. The allowed trait ranges are shown. (C-E) Final trait density distributions (in grayscale black is most dense) and spatial layout. C) With no migration and variation in proliferation we reach 10,000 cells at 26.5 days. D) When proliferation is constant (18 hour intermitotic time) and migration speed varies we reach 10,000 cells at 9.75 days. E) When both vary we reach 10,000 cells at 8.75 days.

## II. Bounding the Trait Space Affects Tumor Growth

In the first case (Fig. 1C), we grow a heterogeneous population of cells with respect to proliferation rate. We start 40 cells with proliferation rates normally distributed around a mean of 18 hours (h) with a standard deviation of 4 h and no migration and grow the mass to 10,000 cells. Unsurprisingly, we find that quickly the fast proliferators take over the proliferating rim, and the slower proliferators are caught in the quiescent core. It takes 26.5 days to grow to this population size. In the second case (Fig. 1D), we grow a population of cells, all with the same proliferation rate (18 h), now with heterogeneity across migration speed (distributed normally around 12 μm/h with a standard deviation of 5 μm/h). The most dominant clones are not so obvious in this case. There are regions with fast migrators and regions with more middle range migrators. The competition amongst cells of different migration speeds is more complex. Incorporating migration disperses the mass and decreases the selection pressure for all. This tumor grew to 10,000 cells in 9.75 days, less than half the time of the tumor with variation only in proliferation. Lastly, we vary both traits (Fig. 1E). We find that, while there is still some heterogeneity, the cells that are both fast proliferators and fast migrators dominate, and this tumor grows the fastest, reaching 10,000 cells in 8.75 days.

Competition from lack of space creates a selective pressure on a diverse population to narrow toward clones with faster proliferation rates and migration speeds. However, these traits do not contribute equally to increasing fitness, especially in a population with a mix of phenotypes. Having diversity in the migration speed of cells benefits all by spreading everything out and allowing even slower proliferators and migrators to divide. It is worth noting that there may be an intrinsic trade-off that would restrict these traits from being maximized simultaneously. These ideas and more are fully developed in [11] along with the effects of spatial distribution and the mode of inheritance of traits.

## III. Quick Guide to the Methods

When it comes to diffusive tumors, the ability for a cell to move off grid affects cell dispersion and therefore local space limitation. The methods presented here show a very simple way to prevent overlapping of cells that are proliferating and migrating. More details on the cellular processes and how they are implemented are given below.

*A. Division*: At each time point for each cell, we check the surrounding space for immediate neighbors, which would lie within a distance of 4 cell radii (r), the farthest possible position in which overlap can occur. We record each neighbor’s angular position with respect to the cell, and exclude the block of angles from a bank of 360 degrees in which the cell cannot divide into due to overlap. The excluded angles (θ) are

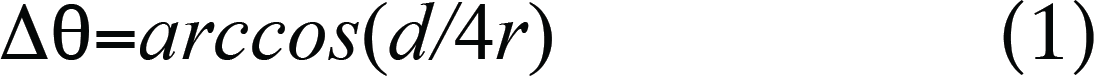

to each side of the line joining the two cells, where d is the distance between cells (as shown in Fig. 1A).

*B. Migration*: The cells follow a persistent random walk. In this case, the persistence times are drawn from a normal distribution of centered around 80 min with a 10 min standard deviation. At each time frame, we identify whether a collision has occurred with another cell, so the time step must be small enough that the fastest moving cells will detect the overlap. At up to 25 μm/h with 1 min time frames, we detect any overlap over ∽0.42 μm. For the collision response, we rewind within the time frame to the moment of contact, then move during the remaining time within the frame with a new persistence and a new angle.

*C. Back to the lattice*: Even when using an off-lattice model, it is occasionally necessary to create a lattice. If, for instance, we want to use a concentration field external to the cell, it will need to be defined within a lattice. As a cell secretes a substance, it can deposit the substance in the lattice point closest to its center, in a few points around the edges, or with a distribution around its center, depending on the lattice size and how detailed one wishes to be. For the cells to respond to the substance, we must define a sampling neighborhood, similar to the deposition neighborhood, as this will regulate the amount of substance the cell senses.

## IV. Alternative Off-Lattice Models

Lattice-based models have many advantages. They are easy to formulate, easy to explain, can be very computationally efficient, and can capture many phenomenological details for most solid tumors. However, these models represent cells as lattice sites, restrict the tissue geometry, and simply look unrealistic. Moreover, velocities are hard to represent, as are more mechanical forces that can occur between cells, which makes the anisotropy associated with invasion and migration harder to achieve.

There are alternative types of off-lattice models that might be better suited for other scenarios [12]. Examples include studies of migration and invasion [13, 14, 15, 16, 17] growing *in vitro* monolayers and spheroids [18, 19], cell adhesion [14], intravasation [20], morphology and patterning [16, 21, 22, 23, 24, 25, 26, 27], and cell/tissue mechanics [23, 24, 26, 27, 28, 29]. These methods can also be used in models outside of cancer, such as embryonic development [30, 31], tissue morphological rearrangement and regeneration [32, 33], and wound healing [34].

When there are no space restrictions, cells can grow exponentially. But in some tumors, when the tissue gets too dense, there may be restrictions from nutrients, growth factors, and space, which can prevent further proliferation and lead too quiescence or necrosis. Considering this, the fraction of proliferating cells will decrease, so the growth curve starts to look more like a power law than an exponential [11]. Some cells can also lose adhesion and pile up more on top of each other. In a tissue, loss of adhesion and piling up could mean more cells will fit in a space and cause less of a space restriction (increasing packing density). Increasing migration or the degree of overlap (decreasing the effective radius) can reduce the effects of this phenomenon and increase the proliferating fraction as shown in Fig. 2.

The method described here is used to study the evolution of a population of cells with regards to how two traits are passed on over generations. However, one must choose the most appropriate model for the scenario in question. Cell migration and spatial restriction are significant concerns in modeling of cancer progression, so working on an off-lattice system can greatly enhance the understanding of invasion, cell interactions, and diverse tumor morphologies. As single cell data becomes more prominent, we can easily incorporate measured cell velocities, persistence times, and turning angles as well as many other cell traits to study heterogeneity through space and time.

**Figure 2.**
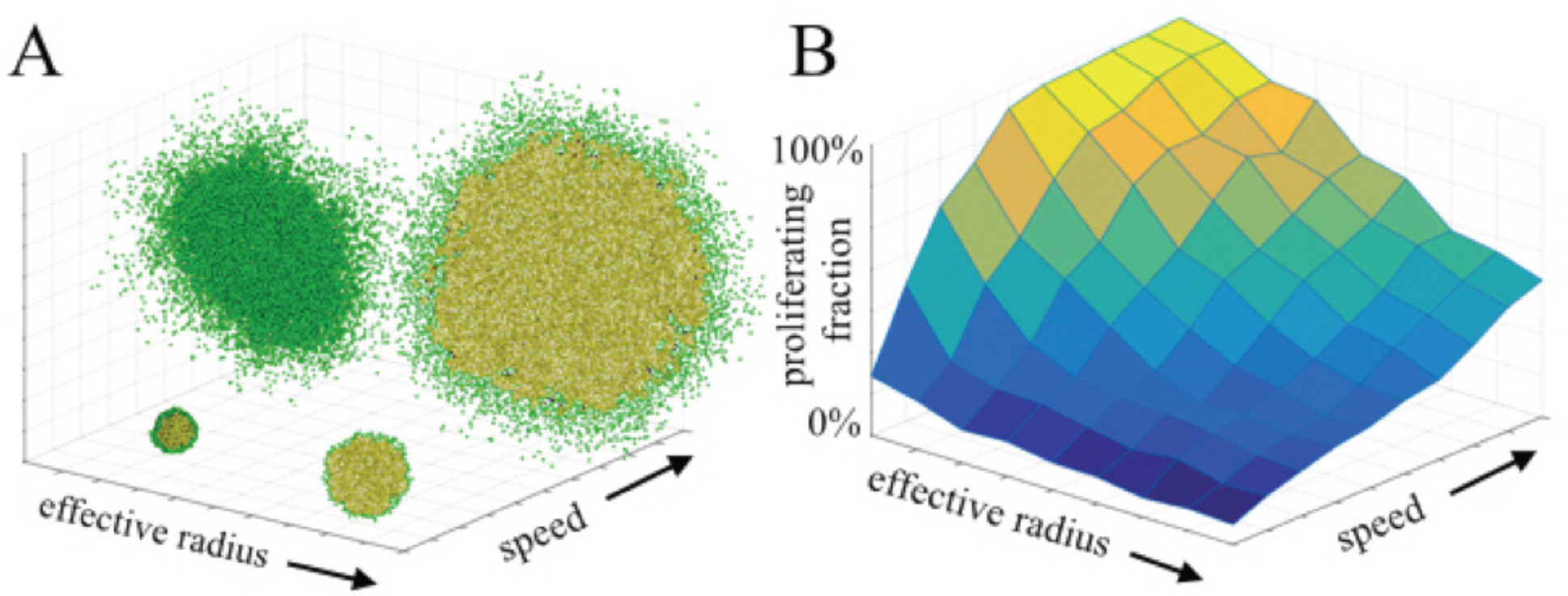
Varying migration speed and effective radius (defines the amount of allowed overlap) affects the population’s proliferating fraction. A) A look at the extreme values for effective radius and migration speed. Green cells are in a proliferating state and yellow cells are in a quiescent state. B) The proliferating fraction increases as the migration speed increases and as the effective radius of a cell reduces.

